# Reticulospinal Tract Hyperexcitability in the Upper Limb After Stroke is Associated with Motor Impairment and Not with Functional Compensation

**DOI:** 10.64898/2026.03.26.714547

**Authors:** Adi Lorber-Haddad, Noy Goldhammer, Tamar Mizrahi, Shirley Handelzalts, Lior Shmuelof

**Affiliations:** Department of Industrial Engineering and Management, Ben-Gurion University of the Negev, Be’er Sheva, Israel; The School of Brain Sciences and Cognition, Ben-Gurion University of the Negev, Be’er Sheva, Israel; The Lillian and David E. Feldman Research Center for Rehabilitation Sciences, Adi Negev - Nahalat Eran Medical Center, Ofakim, Israel; Neurologic Rehabilitation Department at ADI Negev-Nahalat Eran, Ofakim, Israel; Department of Physical Therapy, Ben-Gurion University of the Negev, Be’er Sheva, Israel

**Keywords:** stroke, reticulospinal tract, motor impairments, upper limb function, start react, hyperexcitability

## Abstract

**Background:** Accumulating results suggest that reticulospinal tract (RST) excitability increases after stroke. While animal studies suggest this hyperexcitability may compensate for corticospinal tract (CST) damage, its role in motor function in people with stroke (PwS) remains debated. This study aimed to: (1) replicate findings of RST hyperexcitability in PwS using the StartReact paradigm, measuring acceleration of motor response to a startling auditory stimulus; (2) examine the relationship between RST hyperexcitability and motor impairments after stroke; and (3) explore whether RST hyperexcitability provides functional benefits in severely impaired PwS.

**Methods:** Forty-six PwS completed the StartReact paradigm and motor assessments (Fugl-Meyer, ARAT, grip strength, Modified Ashworth Scale). PwS were categorized into high StartReact effect and typical StartReact effect subgroups based on comparisons with a healthy control group (n=37). Severe impairment was defined as ARAT ≤10.

**Results:** PwS exhibited significantly greater StartReact effects than controls. The high StartReact effect subgroup showed worse motor function, weaker grip strength, and higher spasticity. Among severely impaired PwS, high StartReact effect was not associated with improved grip strength.

**Conclusions:** These findings confirm the existence of RST hyperexcitability after stroke and suggest it is associated with poorer motor outcomes, likely due to reduced cortical input to the brainstem. The absence of functional benefit in severely impaired individuals supports the interpretation that RST hyperexcitability is a maladaptive rather than a compensatory reaction to brain damage. These findings provide insight into the neurophysiological mechanisms underlying motor impairments after stroke and do no imply direct clinical or therapeutic applications.

## Introduction

Six months after stroke, nearly half of persons with stroke (PwS) continue to demonstrate residual motor impairments (Benjamin, 2017). These impairments include negative motor symptoms, such as weakness and impaired selective motor control, largely attributed to injury of the corticospinal tract (CST)(Glover & Baker, 2022; Lawrence et al., 2001; Richardson et al., 2011). The CST provides the primary descending drive for fine and fractionated movements, particularly of distal musculature; consequently, CST damage reduces voluntary activation of spinal motoneurons and leads to decreased strength and impaired motor execution (Daghsen et al., 2024).

Motor deficits also include positive motor symptoms, including spasticity, abnormal synergies, and hyperreflexia. Although traditionally labeled “positive” because they represent excessive motor output, these manifestations have clear functional consequences and are often disabling. These symptoms are thought to arise, in part, from hyperexcitability of the reticulospinal tract (RST). The RST originates in the pontomedullary reticular formation, receives modulatory cortical input, and contributes to posture, proximal limb activation, and gross motor coordination (Glover & Baker, 2022). Following stroke, reduction of cortical inhibition can lead to disinhibition of reticular formation neurons, producing increased RST output (Jang & Lee, 2019; Li et al., 2019; Zaaimi et al., 2012). RST hyperexcitability after stroke has been demonstrated using electrophysiological measures such as ipsilateral motor-evoked potentials (iMEPs) elicited by transcranial magnetic stimulation, which indicate enhanced ipsilateral descending drive consistent with increased RST influence (Mooney et al., 2024).

Behaviorally, heightened RST excitability can be detected through startle-related responses (Kumru & Valls-Solé, 2006; Valls-Solé et al., 2008). The StartReact effect-the shortened reaction time induced by a startling sound-has been linked, in non-human primates, to increased neural activity in the reticular formation (Tapia et al., 2022). In humans, PwS with hemiparesis retain startle-evoked responses despite CST damage, indicating that brainstem pathways mediating rapid motor output, including the RST, remain functionally preserved. Furthermore, PwS demonstrate a larger StartReact effect compared with healthy individuals (Choudhury et al., 2019; Honeycutt & Perreault, 2012). These findings suggest that, in human subjects, following damage to cortical projections, motor control may partially shift toward brainstem-mediated pathways.

The functional contribution of RST hyperexcitability to motor recovery remains unclear. In both stroke and spinal cord injury (SCI), increased RST drive has been associated with severe impairment, including greater expression of pathological synergies and abnormal tone, which can hinder function and response to treatment (Baker & Perez, 2017; Li, 2017; Mooney et al., 2024). Conversely, animal studies demonstrate that recovery following near-complete CST lesions is accompanied by increased reliance on RST pathways (Zaaimi et al., 2012). Similarly, in SCI, RST contributions support gross hand functions, such as power grip (Baker & Perez, 2017).Together, these findings have led to the hypothesis that, in PwS with substantial structural damage to the descending corticospinal fibers-the RST may provide an alternative pathway for motor control (Choudhury et al., 2019). In this context, RST hyperexcitability may simultaneously contribute to maladaptive features (e.g., synergies, tone) while enabling residual gross motor output.

Despite these indirect accounts, direct evidence that RST hyperexcitability supports functional motor performance in humans with stroke is limited. Clarifying whether increased RST activity is primarily functional compensatory, maladaptive, or context-dependent has important implications for prognosis and targeted rehabilitation.

By comparing PwS exhibiting hyperexcitability to those with typical RST excitability, this study examines how RST excitability relates to motor impairment and whether reliance on this pathway may contribute to residual motor function after stroke.

The goals of this study are: To replicate evidence of RST hyperexcitability in PwS using the StartReact paradigm, to characterize the association between RST hyperexcitability and motor impairments, and to explore whether heightened RST excitability confers functional advantage in severely impaired individuals.

## Materials and Methods

The study was conducted at the Adi Negev - Nahalat Eran Rehabilitation Center in collaboration with the Lillian and David E. Feldman Research Center for Rehabilitation Sciences. This study was approved by the Helsinki Ethics Committee of Adi Negev Hospital (Ethics Code: ADINEGEV-2023_106) on 09.12.2023. All participants provided written informed consent prior to enrollment in the study.

### Participants

PwS were recruited from the inpatient neurological rehabilitation department and the outpatient rehabilitation departments at the Adi Negev rehabilitation center. Participants were required to be above 18 years old and had to have experienced a first unilateral stroke, ischemic or hemorrhagic, as verified by CT or MRI scans. Participants were also required to be independent in basic activities of daily living (BADL) prior to the stroke. Additionally, all participants had to be able to provide informed consent and comprehend the tasks they were required to undertake during the study. This capacity was evaluated by their attending clinicians. Participants were excluded if they had a traumatic brain injury, spinal cord injury or a history of physical or neurological conditions that could have affected the study procedures or the evaluation of motor function. This included conditions such as severe arthritis, severe neuropathy, or Parkinson’s disease. Participants who had experienced a cerebellar stroke were also excluded from the study.

The study included 46 PwS (13 females, 33 males; mean age: 65.9 ± 11.7 years) and 37 healthy controls (27 females, 10 males; mean age: 64.8 ± 8.9 years). Participants’ demographic characteristics for the PwS and control groups are summarized in Table 1. No significant difference in the StartReact effect was found between the right and left hand in controls (U = 181, p = 0.55).

**Table 1:**
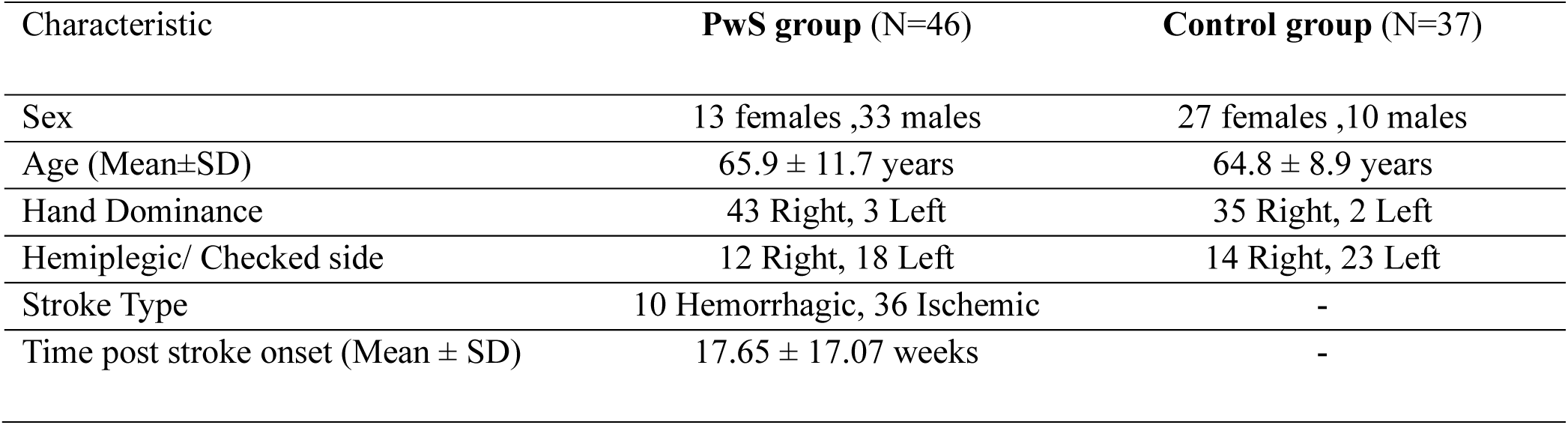
Demographic characteristics of the PwS and Controls.

### Experimental Procedure

Participants in the PwS group underwent functional and motor assessments for the upper limb. We employed the StartReact paradigm to assess RST hyperexcitability, following the approach described in previous studies (Choudhury et al., 2019). Participants completed the clinical and motor assessments, as well as the StartReact paradigm, on the same day.

### Clinical measures

Motor impairment and functional evaluations included 1) the Upper Extremity Fugl-Meyer Assessment (UE_FMA)(Fugl-Meyer et al., 1975; Gladstone et al., 2002), 2) the Action Research Arm Test (ARAT) (Chen et al., 2022; Lyle, 1981), 3) grip strength measured with a microFET dynamometer (Hogrel, 2015; Massy-Westropp et al., 2011), and 4) muscle tone measured using the Modified Ashworth Scale (MAS)(Dunning, 2011; Meseguer-Henarejos et al., 2018; Richard W. Bohannon, 1987) for the wrist extensors (extensor carpi radialis longus, extensor carpi radialis brevis, extensor carpi ulnaris) and elbow flexors (biceps brachii, brachialis, brachioradialis).

The Fugl-Meyer Assessment (FMA) has demonstrated inter- and intra-rater reliability ranging from 0.98 to 1, with a test–retest reliability of 0.96 (Duncan et al., 1983; Mao et al., 2002). FMA impairment levels were defined according to Woytowicz et al. (2017) who classified upper-limb motor severity using cluster analysis of Fugl–Meyer Upper Extremity scores: Severe = 0–28, Moderate = 29–42, and Mild = 43–66. For the Action Research Arm Test (ARAT), inter-and intra-rater reliability values range from 0.96 to 0.99, and test–retest reliability is 0.96 (Platz et al., 2005). Grip strength measured by dynamometry has demonstrated test–retest reliability ranging from 0.947 to 0.967 (Hogrel, 2015). Grip strength measurements were converted to z-scores, adjusted for each participant’s age, sex, and dominant hand (Massy-Westropp et al., 2011). In cases where participants were unable to perform a power grip, the raw grip-strength score was recorded as 0 kg and included in the Z-score calculations. MAS has demonstrated intra-rater reliability ranging from 0.71 to 0.94 in the upper extremities (Vidmar et al., 2023). One potential limitation of MAS is that this measure is not sensitive to the severity of spasticity (Frenkel-Toledo et al., 2021; Piscitelli et al., 2025). Therefore, we used the MAS measure as a binary indication of spasticity (MAS ≥ 1)(Cheng et al., 2023; Schinwelski et al., 2019). In cases of flaccidity, the muscle tone assessment was not performed and therefore excluded, as MAS score of 0 represents normal muscle tone.

The functional assessments were performed by 4 trained OTs and PTs assessors from the Negev laboratory clinical team (Hervé-Colas et al., 2026).

### The StartReact paradigm

Although StartReact responses can be recorded across muscle groups, reticulospinal output is inherently asymmetric, with stronger drive to proximal and flexor muscles, as shown in healthy individuals (Eilfort et al., 2025). Following CST damage, this asymmetry is further amplified, with increased reticulospinal input to flexor muscles and minimal modulation of extensor pathways (Zaaimi et al., 2012), consistent with the characteristic pattern of flexor overactivity and extensor weakness observed after stroke (Choudhury et al., 2019; Lum et al., 2003). Therefore, the assessment targeted both proximal flexors and distal extensor muscles.

The assessment was performed in two consecutive parts. First, on the elbow flexors and second, on the wrist extensors of the more affected side. The StartReact effect was assessed using surface electromyogram (EMG) recordings from electrodes placed over the biceps and the wrist extensors compartment (extensor carpi radialis brevis & longus, extensor digitorum Communis). This selection was guided by the rationale of capturing RST-related activity in movements that considered to be within a flexor synergy (elbow flexion) and outside a synergy (wrist extensors), across both proximal and distal muscle groups. Participants were seated upright with their trunk supported by the chair backrest. When trunk stability was insufficient, the trunk was secured to the backrest using straps. The affected shoulder was in a neutral position alongside the trunk, with the elbow flexed to approximately 70⁰ and the forearm in midpositioned resting on the lap (Germann et al., 2023; Mooney et al., 2024). Participants were instructed to attend to a red light-emitting diode (LED) placed approximately 1 meter in front of them and to make a rapid concentric elbow flexion movement (first session)(Castellote & Valls-Solé, 2015; Eilfort et al., 2025; Eilfort & Filli, 2025; Honeycutt & Perreault, 2014; Lee et al., 2022; Yang et al., 2019) or a wrist extension (second session) when the LED was illuminated (LED on time: 50 ms).

The LED flash was delivered alone [Visual Reaction Time (VRT)] or paired with either a low (∼80 dB, 500 Hz, 50 ms) [Visual Auditory Reaction Time (VART)] or a loud acoustic stimulus (∼110 dB) [Visual Startle Reaction Time (VSRT)]. The acoustic stimuli were delivered through two passive loudspeakers positioned 1 meter in front of the participant at ear level.

Sound intensity was verified prior to data collection using a decibel meter, ensuring consistent delivery of the acoustic stimuli throughout the experiment. These intensities were comparable to those used in previous studies (Choudhury et al., 2019; Fisher et al., 2013; Nonnekes et al., 2014). Prior to data collection, participants were familiarized with the task by performing four practice responses to the LED cue. Subsequently, five loud auditory stimuli were presented without requiring task execution in order to reduce the overt startle response through habituation. These familiarization and habituation trials were not included in the analysis.

The experiment included 20 repetitions of each condition (VRT,VART,VSRT) for each movements’ sessions (elbow flexion and wrist extension), presented in a random order. Each trial lasted 5-seconds. The experiment consisted of 20 repetitions for each stimulus condition (VRT, VART, and VSRT) during two movement sessions (elbow flexion and wrist extension), presented in a randomized order. Surface EMG signals were recorded using a sampling frequency of 2400 Hz. Raw EMG data were first centered by removing the DC offset and then filtered using a 20–450 Hz band-pass Butterworth filter to isolate the physiological range of EMG activity. A 50 Hz notch filter was applied to suppress power-line noise, followed by full-wave rectification of the band-passed signal. To obtain the linear envelope of the EMG, the rectified signal was low-pass filtered with a 25 Hz cutoff frequency, using a Butterworth filter, providing a smooth representation of overall muscle activation. Reaction time was defined as the latency between stimulus onset and the first detected increase in EMG amplitude above baseline, operationalized as exceeding the pre-stimulus mean by more than three standard deviations. This measure was computed automatically for each trial using a custom Python script and was verified manually using a semi-automated graphical interface.

For the StartReact effect, we calculated the difference between VART and VSRT (i.e., VART – VSRT) for each trial for each trial (Choudhury et al., 2019; Germann & Baker, 2021). Each participant’s StartReact effect value was defined as the median of all trial-level StartReact effect measurements.

The StartReact paradigm was conducted on a desktop PC located in the laboratory, using a g.USBamp system by g.tec. Experiments were programmed in MATLAB and Python. All StartReact sessions were conducted by the same clinician (ALH).

### Data analysis

To address the first objective, both the stroke group and the control group were assessed using the StartReact paradigm. For the second objective, participants were stratified into two groups based on their StartReact effect: high StartReact effect group was defined as having a larger StartReact effect than the 90^th^ percentile in the healthy control group i.e, StartReact effect = 40.35ms. Selected positive and negative motor symptoms (including synergy patterns assessed with the UE-FMA, spasticity assessed with the MAS, and grip strength assessed with a dynamometer) were compared between groups.

Finally, to explore whether RST hyperexcitability may serve a functional adaptive role, a subgroup of participants clinically classified as having severe motor impairments [ARAT≤10, n=15 stroke participants (Valladares et al., 2024)] was investigated. Studies have shown that severe CST damage, defined by extensive lesion overlap or absent motor evoked potentials (MEP−), is a major determinant of poor upper-limb motor performance and ARAT scores (Hordacre et al., 2021; Lam et al., 2018). Within this subgroup, we examined whether RST hyperexcitability, reflected by a high StartReact effect, is associated with functional advantages (e.g., greater grip force) compared with individuals who do not exhibit this response, indicating that this neural alternative mechanism may translate into functional compensation.

### Statistical analysis

Prior to conducting group comparisons, data was evaluated to confirm the assumptions of parametric testing. The Shapiro-Wilk test indicated significant deviations from normality across all variables (p < .01), while Levene’s test revealed heterogeneity of variances (p < .01). Non-parametric statistical methods were therefore selected. Group differences were examined using the Mann-Whitney U test, which provides a robust alternative to parametric tests in the presence of non-normal distributions and unequal variances.

In addition to analyzing continuous variables, categorical variables were summarized using contingency tables to explore the distribution of group characteristics. To assess associations between categorical variables (e.g., the presence of spasticity), Chi-square tests of independence were conducted. This approach allowed for the evaluation of whether the observed frequencies significantly deviated from expected distributions under the assumption of independence between variables.

All statistical analyses were performed using JASP statistical software.

## Results

### Differences in StartReact effect between PwS and Controls

Comparison of the StartReact paradigm reaction times between PwS and control groups revealed significantly slower biceps responses in PwS in all three experimental conditions: Visual Reaction Time (VRT)(control median:180.8 ms, PwS median: 231.4 ms, *U* = 483, *p* <.01), Visual Auditory Reaction Time (VART) (control median:164.6 ms, PwS median: 201.1 ms, *U* = 472, *p* < .01), and Visual Startle Reaction Time (VSRT) (control median:163.2 ms, PwS median:167.9 ms, *U* = 601.5, *p* = .01) (Figure1A), indicating deficits in movement initiation, likely to be associated with damage to the CST system.

Additionally, the enhancement in StartReact effect (VART-VSRT difference) was greater in the stroke group [VART-VSRT (PwS median:15 ms, control median: 5.21 ms, *U* = 667.5, *p* = .047 (Figure 1B)], indicating that PwS exhibit increased startle response compared to control subjects (Choudhury et al., 2019; Tapia et al., 2022).

**Figure 1:**
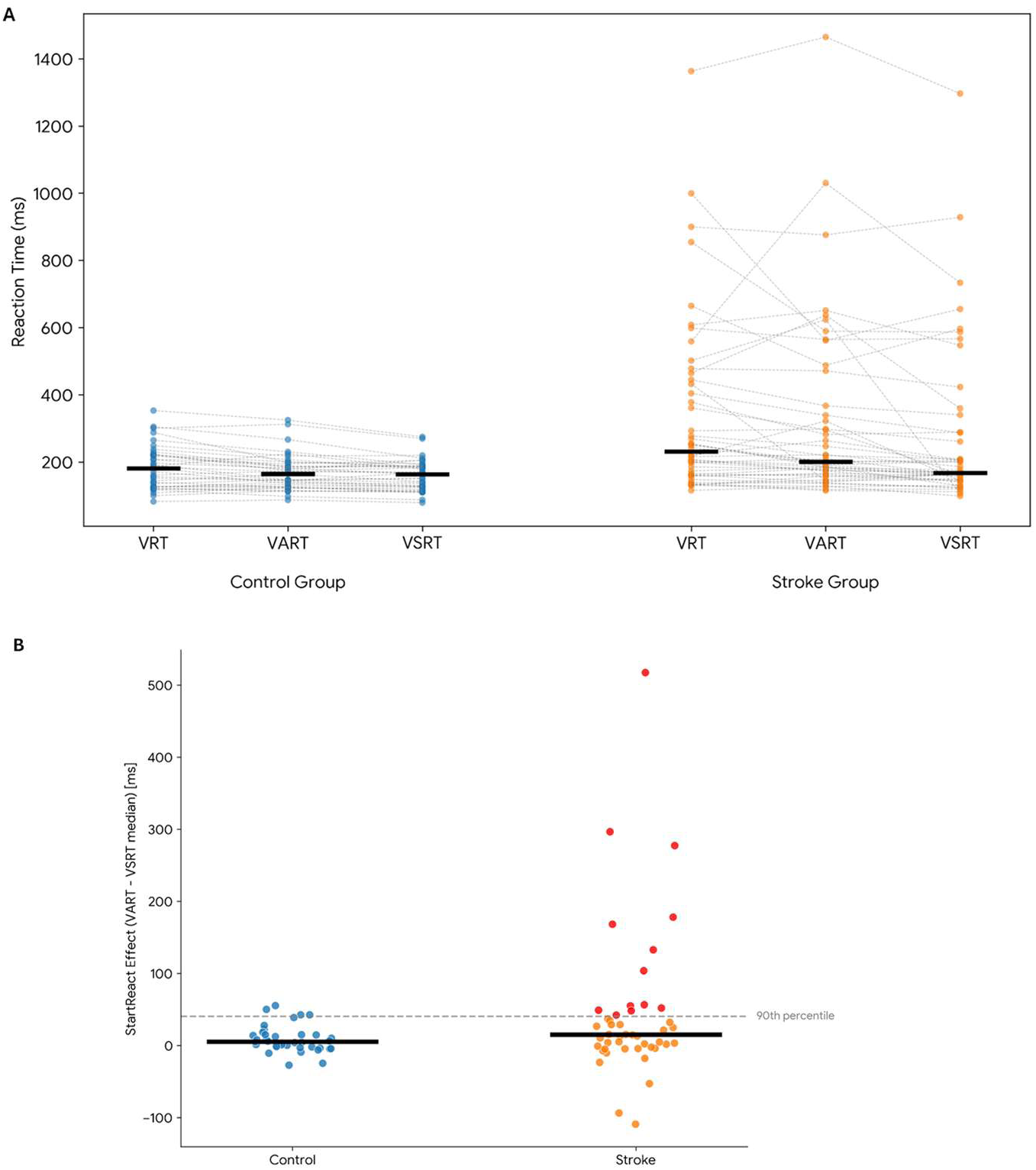
The StartReact effect during elbow flexion in PwS vs. controls. A. Median reaction times (black horizontal lines) in PwS and controls. (each participant is represented by a point) across conditions: Visual Reaction Time (VRT), Visual Auditory Reaction Time (VART), and Visual Startle Reaction Time (VSRT). B. The StartReact effect in PwS and controls during elbow flexion. Participants above the 90^th^ percentile (horizontal dashed line) are shown in red.

The PwS group also presented increased inter-subject variability in the StartReact effect (Figure 1B), with some subjects showing comparable StartReact effect to the control group, and some showing increased StartReact effect with respect to the control group. To further study the phenotype of RST hyperexcitability, we divided the PwS into two groups based on their StartReact effects: subjects with a high StartReact effect and subjects with a typical StartReact effect. The threshold for defining the high StartReact effect sub-group was the 90th percentile of the control group’s reaction time distribution (i.e, StartReact effect = 40.35 ms). PwS with StartReact effect values exceeding this threshold were categorized as having a high StartReact effect, while the remaining subjects were classified as exhibiting a typical StartReact effect.

### Motor Impairment differences in PwS with high StartReact effect and PwS with typical StartReact effect

After confirming that subjects after stroke show high StartReact effect, we turned to study the association between high StartReact effect and motor impairment. Moderate negative correlations were observed between StartReact effect and both clinical measures of upper limb motor function. Specifically, StartReact effect was negatively correlated with ARAT scores (r = –0.44, p < .01) and with FMA scores (r = –0.51, p < .01) (Figures 2 A and B, respectively). The subgroup analysis (e.g high StartReact effect vs. typical) revealed that both the total UE-FMA score and its sub-components (upper extremity, wrist, hand, and total score), were significantly worse in the high StartReact effect subgroup (FMA upper extremity: U=101.5, p<0.01; FMA wrist: U=104.5, p<0.01; FMA hand: U=79.5, p= < .01; FMA total: U=85.5, p<0.01, Table 2).

**Figure 2:**
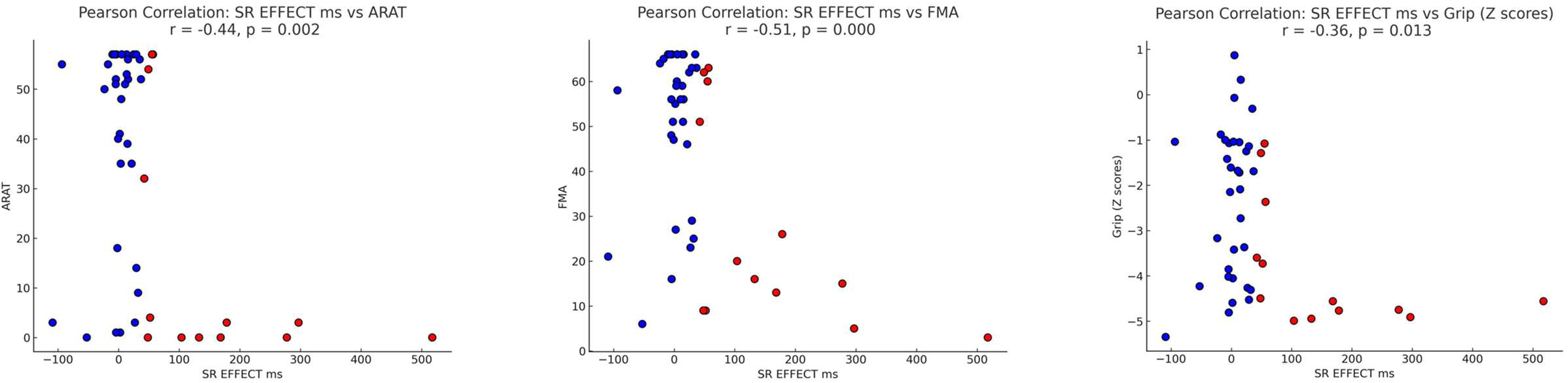
Scatter plot depicting the association between ARAT (A), FMA (B) and GRIP Z scores (C) and StartReact (SR) effect. Participants in the high StartReact subgroup appear in red.

**Table 2:** Clinical Characteristics of PwS by StartReact effect level: Hyper vs. Typical.

|  | High StartReact effect | Typical StartReact effect |
| --- | --- | --- |
| Number of Participants | N=13 | N=33 |
| Stroke Type | 4 Hemorrhagic, 9 Ischemic | 6 Hemorrhagic, 27 Ischemic |
| Phase (Chronic vs. Subacute and Mean weeks after stroke ± SD respectively) | 5 Chronic (38.8± 11.9)<br>8 Sub-acute (4.7±3.1) | 14 Chronic (33.9± 13.5)<br>19 Sub-acute (5.6± 2.6) |
| Number of participants in FMA severity sub-scale (Woytowicz et al., 2017) | 4 mild, 0 moderate, 9 severe | 27, mild, 1, moderate, 5 severe |
| FMA median | 16 | 58 |
| Number of participants in ARAT severity sub-scale (Valladares et al., 2024) | 9 No Activity<br>0 Poor Activity<br>1 Limited Activity<br>1 Notable Activity<br>2 Full Activity | 6 No Activity<br>2 Poor Activity<br>5 Limited Activity<br>8 Notable Activity<br>12 Full Activity |
| ARAT median | 3 | 51 |
| Number of participants in each MAS score (0,1,+1,2,3,4) | 4,4,0,0,4,0 | 27,8,0,1,8,1 |

The ARAT scores were also decreased in the high StartReact effect subgroup (U=103, p<0.01, Table 2). Taken together, individuals with high StartReact effect tended to show greater motor impairments. These results align with previous findings, showing a positive association between high StartReact effect and motor impairment severity (Choudhury et al., 2019; Mooney et al., 2024).

A common symptom after stroke is muscle weakness, which can be assessed through grip force (Olczak, 2021; Riddle, 1989). Maximal grip force depends on proximal shoulder and elbow stability, as well as adequate wrist extensor activation, both of which are commonly impaired after stroke (Ekstrand et al., 2016; Horsley et al., 2016; Mandalidis & O’Brien, 2010). A moderate negative correlation was found between grip strength and the StartReact effect measured in the biceps (r = −0.36, p = 0.01)(Figure 2). Subgroup analysis shows that grip strength is significantly lower in individuals with high StartReact effect (mean Z scores −3.85 in the High StartReact group VS. −2.32 in the Typical StartReact effect group, U= 95, p< 0.01). To summarize, individuals with StartReact effect tend to have weaker grip strength.

One of the functions of the RST is regulating muscle tone (Brown, 1994; Li, 2017). A pathological RST hyperexcitability is therefore suspected to be associated with elevated spasticity levels, as reflected in higher MAS scores (MAS ≥ 1). Indeed, Mann-Whitney U test revealed a significant greater biceps spasticity in the high StartReact effect subgroup (U = 266, p = 0.02). Comparing the differences in probabilities of exhibiting spasticity in the high and typical StartReact effect subgroups, supported the observed association between spasticity and high StartReact effect (61.54% vs 38.46%, χ² = 3.82, p = 0.05).

To test the differential RST excitability across muscle groups, we examined whether StartReact effect extends to movements outside the flexor synergy, by examining StartReact effect in wrist extensors. Among the 46 stroke subjects, 28 demonstrate discernible wrist extension EMG activity. Within this subgroup, no significant differences were observed between the PwS and the control group (VART-VSRT: control median: −0.36 ms, PwS median: 5.56ms, U = 339.00, p = 0.39). Furthermore, PwS with high StartReact effect in the wrist extensors (i.e, SR effect ≥ 11.98ms.) did not show increased impairment according to their ARAT and FMA scores. ARAT (U = 109.5, p = .46), total Fugl-Meyer score (FM sum: U = 112.5, p = .38), Fugl-Meyer upper extremity (FM upper extremity: U = 109, p = 0.47), Fugl-Meyer wrist subscore (FM wrist: U = 89.50, p = 0.85), or Fugl-Meyer hand subscore (FM hand: U = 106, p = 0.54). Last, no correlation was observed between the magnitude of the proximal (biceps) and distal (wrist) StartReact effects (r = –0.11, p = 0.57), nor between the StartReact effect and the level of motor impairment as measured by the FMA (r = −0.09, p = 0.63) and ARAT (r = −0.17, p = 0.39).

Taking together, the StartReact effect in the wrist extensors was not associated with motor impairments and was not affected by stroke, supporting the weaker contribution of RST to these muscles.

### The association between time after stroke and the StartReact effect

Our population consisted of participants both in the chronic and in the sub-acute phases (Mean weeks after stroke 35.2 ± 12.9 SD, 5.3± 2.7 SD respectively). This variability may provide information about possible changes in RST excitability as a function of time after stroke. RST excitability did not show a significant difference as a function of time post-stroke [Chronic StartReact effect median= 21.46 ms, Sub acute StartReact effect median= 13.33ms, U= 0.60, p= 0.55]. Additionally, the proportion of participants with high StartReact was not significantly different between the two phases [chornicHigh StartReact effect ratio= 5/13 vs. subacute High StartReact effect ratio = 8/13, χ²= 0.06 p=0.81]. Importantly, impairment levels also did not differ between groups [ARAT chronic median: 41.00: sub-acute median: 51; U=217, p=0.38.

FMA chronic median: 55 sub-acute median: 56; U=252, p=0.93. Grip z scores: chronic median: −3.42, sub-acute median: −2.09; U=208.5, p=0.29. spasticity: MAS≥1 chronic: 9/19, sub-acute: 9/27, χ²= 0.92, p=0.34]. We suggest that the lack of differences between groups reflects the specific sampling characteristics of the current study.

### Search for a functional advantage of high StartReact effect in severely impaired PwS

To explore whether RST hyperexcitability has a functional compensatory role, as was suggested by findings from primate studies (Zaaimi et al., 2012), we examined PwS with severe impairment [ARAT score below 10 indicating no activity (Valladares et al., 2024)] which demonstrate discernible EMG activity (N=15) and asked, within this subgroup, if the grip strength is increased in PwS with high StartReact effect. Contrary to our hypothesis, participants with high StartReact effect showed reduced grip strength (median Z score of −4.75 in the high StartReact effect subjects (N=9), vs. −4.29 in the typical participants (N=6)). The Mann-Whitney U showed no significant difference in grip strength between the two groups (*U* =19.00, *p* = 0.38).

Another approach to examine if high StartReact effect is associated with a functional advantage is to compare the proportion of participants with high StartReact effect across the range of motor impairment severity (Table 2). If high StartReact effect is associated with a *functional* benefit, we expect to find a reduced proportion of high StartReact effect participants in the most severe side of the range. A chi-square test revealed a significant association between high StartReact effect, and severity of motor impairment based on FMA scores, classified as Mild, Moderate, and Severe according to Woytowicz et al. (2017) (χ²=12.96, p < 0.01).

Specifically, the group with severe impairment showed a higher frequency of high StartReact effect participants (9 out of 13), while the mild impairment group showed a predominance of typical StartReact effect participants (27 out of 31). When running a similar analysis based on the ARAT score-focusing on the participants with ARAT≤10, we also did not find a reduced proportion of the high StartReact effect participants. On the opposite - a significantly higher proportion of high StartReact subgroup (60% compared to 28.2% in the total group) was found (p < 0.01), indicating high StartReact effect in this group of low ARAT scores. The consistent findings across the two analyses suggest that high StartReact effect is more common in individuals with greater motor impairment and show no evidence for a *functional* benefits that are associated with high StartReact effects.

## Discussion

### High StartReact effect as an indication of RST hyperexcitability after stroke

As expected, PwS exhibited significantly greater reaction time enhancement (StartReact effects) compared to controls, suggesting RST hyperexcitability after stroke (Tapia et al., 2022). Our findings are consistent with previous studies showing longer StartReact effects in PwS (Choudhury et al., 2019). Furthermore, to study the association between motor impairment and RST hyperexcitability, we divided the PwS to hyperexcitability and a typical excitability subgroup. This approach allowed us to examine the additive effect of RST hyperexcitability on motor impairments after stroke. This approach is supported by a recent study showing that PwS with severe CST damage have higher iMEP signals in the paretic biceps brachii compared to age-matched healthy controls, whereas PwS with mild CST damage (measured by preserved cMEPs), do not differ significantly from healthy controls in their iMEP responses (Mooney et al., 2024).

The absence of significant differences in StartReact effect in wrist extensors between PwS and controls, together with its lack of association with motor impairments, suggests that RST hyperexcitability is manifested primarily in proximal flexor muscles. Indeed, previous studies (Honeycutt et al., 2015; Honeycutt & Perreault, 2014) found increased startle responses in PwS which were biased to elbow flexor muscles (compared to extensors). Consistent with this pattern, studies have shown that although startle responses can be elicited during wrist or finger extension after stroke, their probability is relatively low and they do not differ significantly between PwS and healthy controls. Moreover, reaction times triggered by startling stimuli in this distal extensors are often comparable to those of controls and are not significantly associated with motor impairment severity (Honeycutt et al., 2015). These findings suggest that the RST plays a stronger role in activating proximal flexor muscles, and supports the contribution of hyperexcitability to abnormal motor patterns after stroke, such as flexor synergies (Twitchell, 1951).

### RST hyperexcitability and motor impairments

Examining the potential link between RST hyperexcitability and motor impairments in our study showed that individuals with RST hyperexcitability, as reflected by high StartReact effect, showed more severe motor impairments, and a greater likelihood of spasticity. Our results are consistent with previous studies that shown that PwS with high StartReact effect expressed greater levels of spasticity (Choudhury et al., 2019). Moreover, recent studies describe the role of the RST in the development of spasticity, highlighting that muscle tone is normally maintained by inhibitory input from the CST and facilitatory input from the medial RST (Mukherjee & Chakravarty, 2010). Damage to the CST may reduce dorsal RST inhibition, resulting in disinhibition of stretch and flexor reflexes. Consequently, medial RST activity becomes unopposed, leading to increased muscle tone and spastic posture.

Besides the manifestation of spasticity, severe motor impairments were also associated with high StartReact effect. This finding is consistent with previous research reporting a negative correlation between iMEP presence and upper limb motor function, as measured by FMA (Mooney et al., 2024). Specifically, individuals with lower FMA scores exhibited higher iMEP signals only in response to startling stimuli supporting the role of RST in this modulation.

Our results are in line with previous findings demonstrating that increased StartReact effect in PwS is correlated with ARAT, FMA and MAS scores, indicating greater motor impairment (Choudhury et al., 2019; Mooney et al., 2024). Introducing an objective cutoff-based classification of heightened StartReact effect, allowed us to confirm the association between RST hyperexcitability and motor impairment and a more comprehensive characterization of the behavioral implications of RST hyperexcitability post stroke. Notably, our cutoff-based approach for defining heightened RST excitability do not support previous conjecture that heightened RST outputs may aid recovery and have a functional advantage in severely affected individuals (Choudhury et al., 2019).

### No evidence of functional contribution from RST hyperexcitability

The last aim of our study was to assess whether RST hyperexcitability provides functional benefits in severely impaired PwS. Animal studies have proposed a compensatory role for the RST system following CST damage. For instance, research in macaques demonstrated that the StartReact effect emerged only when brainstem inputs dominated motoneuron drive, highlighting the RST’s capacity to facilitate rapid movement (Tapia et al., 2022). Moreover, CST lesion studies in macaques have suggested that RST excitability may support recovery of gross motor function (Zaaimi et al., 2012), by showing that following unilateral CST lesions, synaptic input from medial brainstem pathways, likely including the RST to forearm flexors, was significantly strengthened. This reorganization supported partial recovery but also introduced a flexor-dominant bias, potentially contributing to abnormal synergies. Similarly, upregulation of cortical inputs to the medial RST in monkeys is strongly associated with improved hand motor recovery especially in reaching and grasping tasks (Darling et al., 2018).

Importantly, the only indication of a possible compensatory function of RST hyperexcitability in humans comes from Spinal Cord Injury (SCI). Using the StartReact paradigm, in individuals with incomplete cervical SCI, a startling cue selectively shortened reaction times during power grip but not during precision grip or finger abduction (Baker & Perez, 2017). Those findings suggest that after SCI, RST supports gross motor functions such as power grip as expected by the physiology of this pathway. Following the same line, RST inputs were shown to be increased to the biceps (flexors) compared to the triceps (extensors), consistent with the flexor bias that characterizes the RST system (Sangari & Perez, 2020). Taken together, these findings raise the question of whether a similar phenomenon is present in stroke.

The present findings demonstrate that RST hyperexcitability is associated with worse motor outcomes, including reduced grip strength and greater impairment severity, without evidence of functional compensation, even among individuals with severe deficits. This pattern suggests that, in humans, RST hyperexcitability is more likely to reflect maladaptive motor response to the neural damage, contributing to phenomena such as spasticity and abnormal synergies, rather than facilitating recovery of grip force, and voluntary movement. These results support previous perspective arguing that the compensatory role of the RST in motor recovery is largely confined to animal studies (Li et al., 2019).

The difference between the compensatory role of the RST among monkeys and humans may be the result of structural organization differences between the species. Specifically, while the CST is highly developed in humans and critical for fine voluntary motor control, in monkeys CST connections are predominantly polysynaptic, and may have more redundancy with other pathways during skilled tasks. Moreover, the RST has denser projections in monkeys, reflecting its prominent role in trunk and limb control (Obara, 2023). In monkeys, the RST demonstrably maintains gross motor function post-CST damage, and its additional lesion severely impairs these skills, with some recovery potentially mediated by other tracts like the rubrospinal tract, which is more prominent in monkeys (Atkinson et al., 2022).

In addition to the reaction of the RST to the decrease in cortical inputs, RST excitability may also be modulated by training. Recent evidence has demonstrated that long-term paired associative stimulation enhances StartReact effect and subcortical excitability, providing indirect support for the plastic potential of the RST. (Germann & Baker, 2021). In addition, following the observation that weight support reduces postural and movements biases, a conceptual model has been proposed in which, in healthy individuals, the CST moderates the RST, maintaining a balance between movement and posture control (Hadjiosif et al., 2024). After stroke, damage to the CST reduces its moderating influence on the RST, which, along with RST hyperexcitability, leads to abnormal resting postures that can “spill over” into abnormal movements. Taken together, while we could not see any clinical benefit of RST hyperexcitability, the RST may be an effective target for modulation in rehabilitation. Further studies are needed to explore the effect of training on modulation of RST excitability and its cortical projections after brain damage.

Several limitations of the present study should be acknowledged. First, the observational design limits causal interpretations regarding the effect of RST hyperexcitability on motor impairment. Additionally, although subgroup analyses, particularly those focusing on individuals with severe impairment, offered important preliminary insights, the generalization of the findings, especially in terms of the prevalence of reticulospinal hyperexcitability across different clinical profiles, and its potential functional contribution is limited by the small number of participants in these categories. These limitations underscore the need for caution in interpreting the present results. Third, while the StartReact effect is a valuable proxy for assessing RST hyperexcitability, it remains an indirect measure and may be influenced by additional factors such as attention, arousal, or task anticipation. Fourth, the PwS participants included both participants at the chronic and sub-acute phases. Assuming a sequential recovery process where positive signs emerged and decreased during the sub-acute phase (Twitchell, 1951), this wide sampling may increase the variability in the results. Additionally, cross-sectional comparisons of sub-acute and chronic groups are prone to sampling biases. The participants in the study were recruited at different stages after stroke, and therefore could not be stratified based on impairment or recovery scores. Longitudinal studies are necessary to clarify whether RST excitability evolves during recovery or remains static. Finally, the current sample was not stratified by lesion location or CST integrity, which may have introduced variability and limited our ability to detect more nuanced associations between RST hyperexcitability and motor outcomes.

Overall, this study provides supportive evidence that RST hyperexcitability, as measured through the StartReact effect after stroke, is linked to both impaired voluntary motor control and increased spasticity. Further analyses highlight a maladaptive rather than compensatory role for RST activity in PwS, underscoring the importance of attenuating the excitability of the RST after stroke.

## Acknowledgments

The study was supported by “*The Lillian and David E. Feldman Research Fund”.* We are grateful to the participants and their families for their willingness to take part in this study, and we extend our appreciation to the medical and research teams at Adi Negev–Nahalat Eran for their support and assistance throughout the process.

## Author Contributions

*Adi Lorber-Haddad:* Data curation; Formal analysis; Investigation; Methodology; Visualization; Writing-original draft; Writing- review & editing.

*Noy Goldhammer:* Software; Project administration

*Tamar Mizrahi:* Project administration

*Shirley Handelzalts:* Writing- review & editing

*Lior Shmuelof:* Conceptualization; Funding acquisition; Investigation; Resources; Supervision;

Writing—original draft; Writing —review & editing.

## Ethics approval statements

This study was approved by the Helsinki Ethics Committee of Adi Negev Hospital (Ethics Code: ADINEGEV-2023_106) on 09,12,2023. All participants provided written informed consent prior to enrollment in the study.

